# Comparative analysis of multiplexed in situ gene expression profiling technologies

**DOI:** 10.1101/2024.01.11.575135

**Authors:** Austin Hartman, Rahul Satija

**Affiliations:** New York Genome Center, New York City, NY, USA; Center for Genomics and Systems Biology, New York University, New York City, NY, USA

## Abstract

The burgeoning interest in in situ multiplexed gene expression profiling technologies has opened new avenues for understanding cellular behavior and interactions. In this study, we present a comparative benchmark analysis of six in situ gene expression profiling methods, including both commercially available and academically developed methods, using publicly accessible mouse brain datasets. We find that standard sensitivity metrics, such as the number of unique molecules detected per cell, are not directly comparable across datasets due to substantial differences in the incidence of off-target molecular artifacts impacting specificity. To address these challenges, we explored various potential sources of molecular artifacts, developed novel metrics to control for them, and utilized these metrics to evaluate and compare different in situ technologies. Finally, we demonstrate how molecular false positives can seriously confound spatially-aware differential expression analysis, requiring caution in the interpretation of downstream results. Our analysis provides guidance for the selection, processing, and interpretation of in situ spatial technologies.

## Introduction

The advent of highly multiplexed, spatially-resolved gene expression profiling technologies promises to substantially enhance our understanding of complex biological systems. By directly capturing the spatial location of molecules and cells, which is lost using single cell RNA-seq technologies, in situ profiling can provide new insights into the interplay between different cell types, the functional organization of tissues, and the influence of the microenvironment on cellular behavior^1–3^. The ability to infer relationships across multiple scales, from molecules to cells to organ systems, represents a powerful advance and has given rise to a wide diversity of academic and commercial profiling technologies to address this need^4–17^. These technologies can be broadly grouped into sequence-based methods which return transcriptome-wide profiles for individual voxels^18–22^, and in situ imaging technologies which perform highly multiplexed hybridization-based profiling with targeted probes.

While both types of approaches exhibit distinct strengths and weaknesses, imaging-based technologies exhibit clear advantages for high-resolution molecular profiling. The use of uniquely barcoded and fluorescent probes enables the multiplexed identification of single molecules, enabling researchers to study both cellular and sub-cellular localization patterns^23^. Moreover, in contrast to sequencing based technologies that may blend the profiles of adjacent cells located within the same voxel, imaging-based technologies can, in principle, unambiguously assign molecules to single cells. Therefore, in situ techniques are uniquely suited to explore the effect of a cell’s interactions and micro environment on its molecular state, and for example, to quantify how cells of a similar type vary in their gene expression patterns at distinct anatomical sites^24–26^.

The burgeoning interest in multiplexed in situ imaging technologies for spatially resolved gene expression profiling has spurred the development of an exciting variety of academic technologies and commercial offerings. These approaches vary not only in the size and composition of their target gene panel, but also in their approaches for signal amplification, detection, and error correction. Key downstream data processing steps, including cell segmentation can also vary across datasets^27–29^. These experimental and computational differences can cause substantial variation in data quality, necessitating a systematic approach to benchmark and compare in situ technologies. Comparative benchmarking has been invaluable for researchers to select among competing single-cell RNA sequencing (scRNA-seq) technologies^30–32^, but similar efforts and benchmarking strategies are still in the process of being identified for imaging-based spatial technologies^33,34^.

Here, we perform a comparative benchmarking analysis of six different in situ gene expression profiling technologies using publicly available datasets of the mouse brain. We find that standard metrics, such as the number of unique molecules detected per cell (i.e. sensitivity), cannot be directly compared across datasets due to substantial differences in panel composition and off-target molecular artifacts (i.e. specificity). We explore multiple possible sources of molecular artifacts, develop new metrics that control for these phenomena, and utilize these metrics to evaluate and compare in situ datasets. Finally, we demonstrate how even relatively minor decreases in molecular specificity can seriously confound spatially-aware differential expression analysis, thereby requiring caution in the interpretation of downstream results.

## Results

The rapid evolution of imaging-based in situ spatial technologies has led to the creation of a diverse set of technologies for multiplexed gene expression profiling. Similar to previous comparative benchmarking approaches for scRNA-seq^30–32^, which highlighted substantial differences in throughput and sensitivity across multiple technologies, benchmarking in situ technologies can help guide users in choosing a technology. Moreover, while many scRNA-seq technologies are often benchmarked using their performance on peripheral blood mononuclear cells (PBMC), in situ spatial approaches typically demonstrate their performance using sections of the mouse brain^35^, as this tissue has well-characterized and complex spatial organization, and exhibits highly conserved structure across animals.

Based on publicly available data in mid-2023, we identified datasets featuring full mouse brain sections (including coronal or sagittal sections), generated with six different technologies (Figure 1a, Supplementary Methods). Our selection of datasets represented publicly released datasets from three commercial technologies: Xenium (10x Genomics), MERSCOPE (Vizgen), and Molecular Cartography (Resolve Biosciences), as well as three published datasets generated in academic labs using MERFISH^13^, STARmap PLUS^14^, and EEL FISH^9^. Our intention was to enable a fair comparison after the authors had the chance to best showcase their technology. Therefore, in our initial analyses we used the outputs provided directly by the dataset generators. Output files for all technologies include gene-cell expression matrices and x-y coordinates denoting the center of each segmented cell. Molecular Cartography, MERSCOPE, Xenium, MERFISH, and EEL FISH additionally provide the exact position of each detected molecule and MERSCOPE, Xenium, and Molecular Cartography provide vertices denoting the segmentation boundary of each cell.

**Figure 1.**
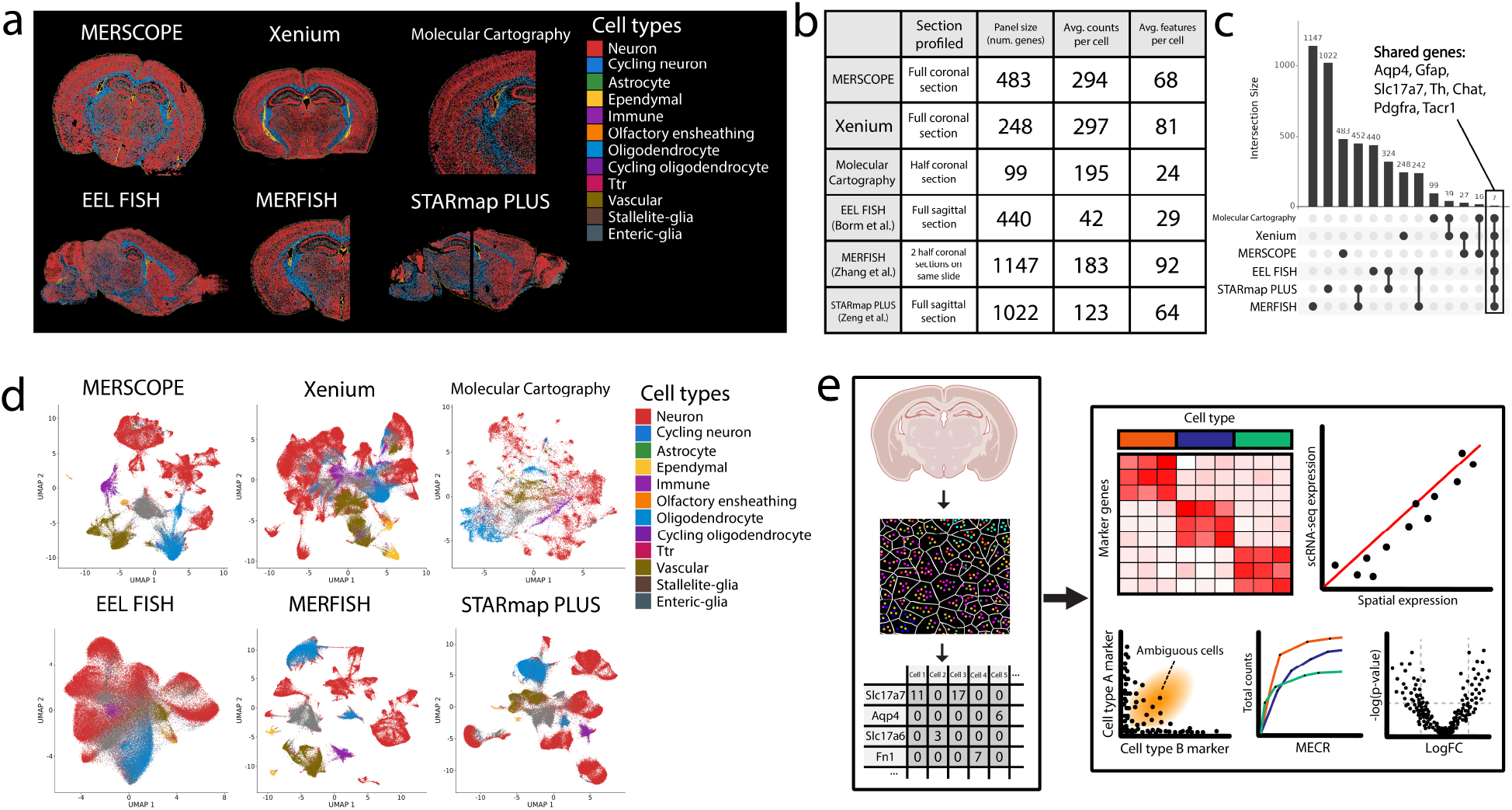
Comparative analysis of in situ gene expression profiling technologies. (**a)** Mouse brain sections from six publicly available datasets. Cells are colored by broad annotation classes derived from scRNA-seq label transfer (Supplementary Methods). **(b)** Overview of technical quality control metrics for each of the six datasets. **(c)** UpSet plot showing the overlap of targeted genes across methods. Seven genes were included in the target panel for all six methods. **(d)** UMAP visualization of each dataset, cells are colored by broad annotation class as in **(a)**. **(e)** Schematic overview of comparative analysis. (Left) We obtained molecular spot calls, cell segmentations and gene expression quantifications from the original dataset generators. (Right) we utilized these data to assess sensitivity, specificity, cell type identification, and spatial differential expression analysis.

We first computed metrics that are commonly reported for scRNA-seq, including the reproducibility across replicate experiments (Supplementary Figure 1a), and the average number of counts and features identified in each cell (Figure 1b). We obtained replicate datasets for five out of the six technologies, and reassuringly observed high reproducibility of pseudobulk values (Pearson correlation ranging from 0.894-0.999) in all cases (Supplementary Figure 1a). Interpreting sensitivity is more challenging, as the panel sizes range substantially across technologies, from 99 genes (Molecular Cartography) to 1,147 genes (MERFISH), with only seven genes being uniformly targeted across datasets (Figure 1c). We observed a range in the average number of total transcripts detected per cell (ranging from 42 for EEL FISH to 297 from Xenium), but observed no correlation between this metric and total panel size (Supplementary Figure 1b). We found that the distribution of average expression levels (based on scRNA-seq data^36^) for targeted genes was similar across technologies, though we did observe an enrichment for very lowly expressed genes (such as G coupled protein receptors) in the MERSCOPE target list (Supplementary Figure 1c).

We annotated each imaged cell from the six in situ datasets based on an adult mouse brain scRNA-seq reference^36^ (Figure 1a,d; Supplementary Methods). We found that all in situ methods were successfully able to delineate broad groups of cells (i.e. oligodendrocytes and astrocytes), and that these annotations were supported by the expression of canonical marker genes (Figure 2a). However, we also observed non-specific expression of these marker genes in other cell types (Figure 2a), particularly in comparison to scRNA-seq (Supplementary Figure 2a). We annotated higher-resolution subsets as well (Supplementary Figure 3a), but we chose not to benchmark annotations given the difference across technologies in panel size and composition. For example, the 10x Xenium dataset includes canonical markers for each of the mouse cortical inhibitory interneuron subtypes (Pvalb, Sst, Lamp5, Vip)^37^, but none of these markers is profiled in the receptor-focused MERSCOPE dataset.

**Figure 2.**
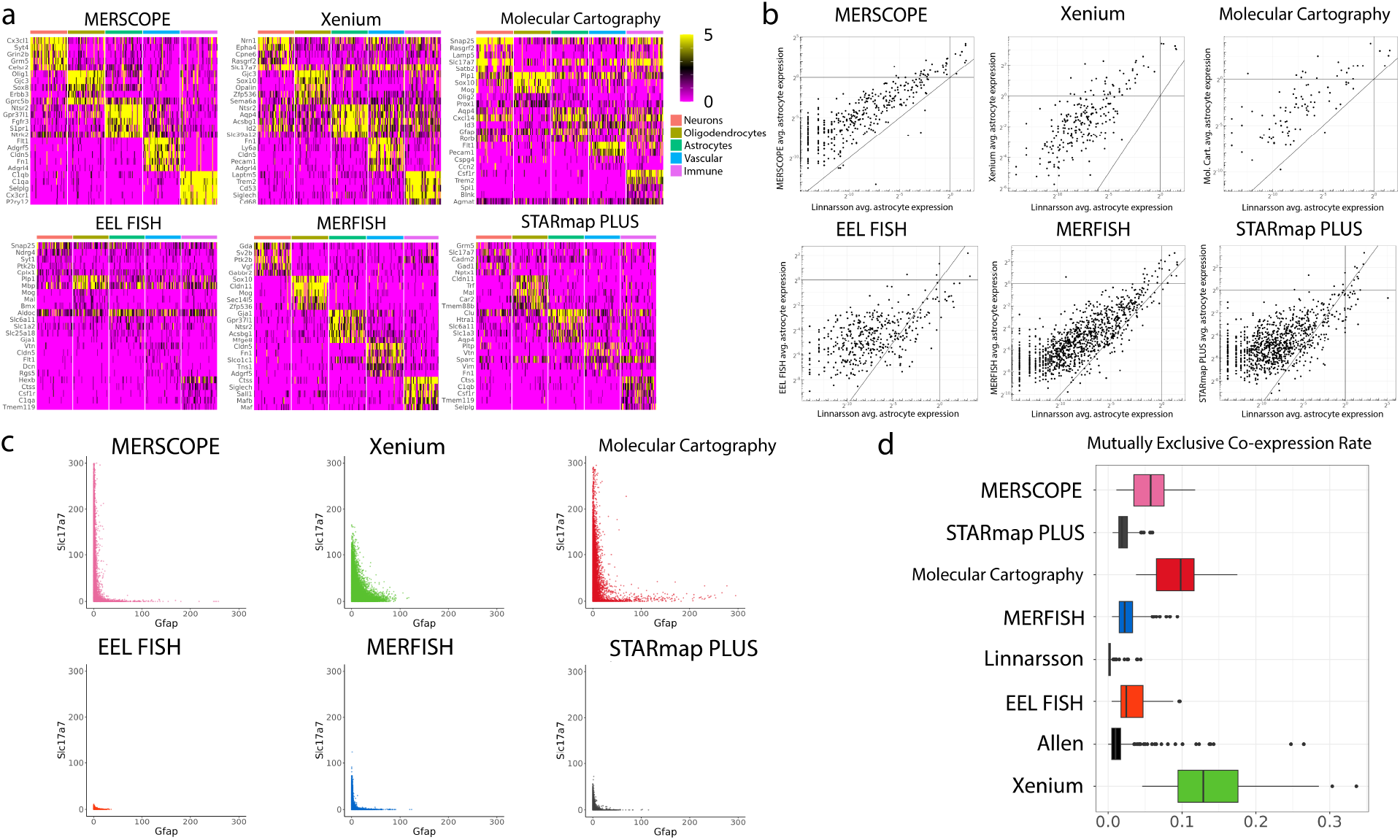
Variation in specificity across in situ technologies. **(a)** Heatmaps displaying marker gene expression (Supplementary Methods) for each of five major cell classes. Heatmap shows 50 randomly sampled cells per class, with a threshold of five molecular counts for each gene. While most technologies exhibit robust marker gene expression in the correct class, we also observe evidence for non-specific expression. **(b)** Pseudobulk expression plots comparing average molecular counts (within astrocytes) for each in situ technology, compared with an scRNA-seq reference. **(c)** ‘Barnyard’ plots showing the expression (counts) of two mutually exclusive genes, Slc17a7 (excitatory neuron marker) and Gfap (astrocyte marker) in each dataset. **(d)** MECR for all six in situ methods. Boxplots display the range of MECR values for all pairs of mutually exclusive genes in each dataset (Supplementary Methods). For reference, scRNA-seq datasets generated using 10x Genomics (Linnarsson) and SMART-Seq2 (Allen) are also shown (Supplementary Figure 4b).

We next performed celltype-matched comparisons of molecular sensitivity between each spatial technology and scRNA-seq. This approach allowed us to assess the sensitivity of the different technologies on a common ground, despite differences in their target panels. In all cases, we found broad quantitative agreement between cell type-specific average single-cell expression profiles computed from in situ and scRNA-seq experiments (Figure 2b). In all six cases, average expression profiles from in situ methods exhibited higher molecular counts for the same gene, when compared to scRNA-seq data. While this finding is consistent with a general assumption that imaging-based technologies can achieve higher sensitivity, it may also be explained in some cases by the presence of non-specific signals which would inflate gene expression counts in imaging datasets. We observed that the relative increase in molecular counts for in situ techniques (compared to scRNA-seq) was highest for lowly expressed genes, but was diminished for highly expressed genes (Figure 2b). This pattern is consistent with the presence of imaging-specific background that has a uniform effect on all probes, and therefore has a proportionally greater effect on probes that return a low number of bona fide molecular counts.

The presence of background RNA has also been observed in scRNA-seq experiments, albeit typically at low levels^38^. The gold-standard test to quantify non-specificity in scRNA-seq datasets is a ‘barnyard’ experiment (i.e. mixing of human mouse cells), where false-positive molecules can be unambiguously identified^39^. While none of the in situ technologies provide a multi-species dataset, we performed a barnyard-style analysis by first identifying genes that exhibit mutually exclusive expression in scRNA-seq data. For example, we found that Slc17a7 and Gfap were specific markers of excitatory neurons and astrocytes respectively, and almost never co-expressed in the same cell (Supplementary Figure 2b). These genes were measured in all six datasets, and while we observed anti-correlated expression in all cases, a subset of technologies (in particular, Xenium and Molecular Cartography) exhibited widespread co-expression (Figure 2c).

To quantify this phenomenon, we calculated a ‘mutually exclusive co-expression rate’ (MECR) for each dataset. This metric aggregates pairs of mutually exclusive genes between cell types based on scRNA-seq and quantifies their rate of detected co-expression, normalized for the abundance of each individual gene (Supplementary Methods). We observe wide variation in this metric across technologies, with the Xenium dataset exhibiting the highest rate across datasets (Figure 2d). We note that based on standard metrics, Xenium exhibits the highest number of average molecular counts per cell (Figure 1b), but that a component of the Xenium molecular profiles represent false positive counts (Figure 2a,c,d). More generally, these results highlight how sensitivity-based comparisons of multiple technologies may be misleading if there are also substantial differences in specificity rate.

We next considered possible sources of non-specific signal for the in situ datasets, starting with the possibility of off-target probe binding or false positive detection. Five of the six in situ datasets also include negative control probes in their binding. These probes represent either probes that are not expected to bind to any human RNA sequence, probes that lack the ability to be successfully converted into fluorescent signals, or both. All datasets exhibited non-specific detection of negative control probes, but typically at levels (Supplementary Figure 3b) that were far below the level of signal observed for bona fide expressed genes. We observe that the ratio of mean counts per gene divided by mean counts per negative probe ranges from approximately 0.001 to 0.1 (Supplementary Figure 3c).

We next considered an alternative factor that can drive dataset-differences in specificity: the accurate assignment of individual molecules to the correct cell. The accuracy of molecular assignment is dependent on the quality and size of cell segmentations, which we found to vary substantially across technologies. For example, we found that the segmentations for the MERSCOPE dataset covered significantly less of the imaged area compared to the Xenium segmentations (Figure 3a,b). This variability is reflective of the inherent challenges in performing cell segmentation and assigning molecules. Overly conservative segmentation maintains high molecular specificity but will discard a substantial fraction of transcripts, while aggressive segmentations may retain a higher fraction of transcripts yet suffer from non-specific assignment. We found that the fraction of molecules assigned to cells, which reflects each dataset’s position on a scale ranging from aggressive to conservative segmentation approaches, was strongly associated with MECR (Figure 3c). We conclude that the choice of segmentation thresholding is important in controlling the balance between sensitivity and specificity for in situ datasets.

**Figure 3.**
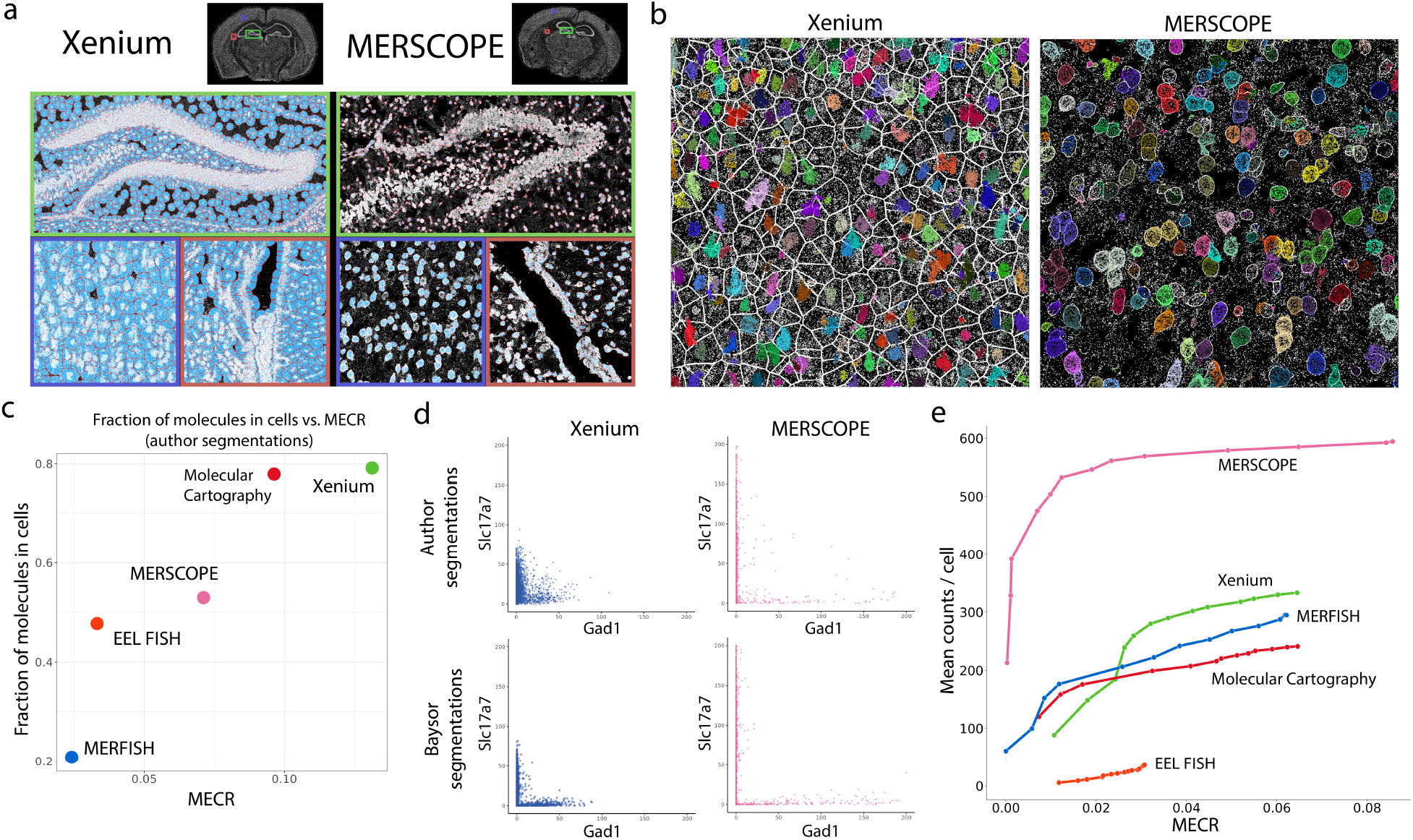
Segmentation size and quality affects molecular sensitivity and specificity. **(a)** Author-provided cell segmentations for the Xenium and MERSCOPE datasets, as representative examples. In these plots, cell borders are colored red, cell interiors are filled in blue, and all detected molecules are overlaid in white. Three representative regions are shown; The dentate gyrus (outlined in green), lateral ventricle (outlined in orange), and a section of the cerebral cortex (outlined in blue). **(b)** Zoomed in regions from the cerebral cortex. Cell boundaries (author-provided) are shown in white. Molecular assignments are provided by Baysor, and are colored by the cell they are assigned to. Unassigned molecules are shown in white. **(c)** Relationship between dataset MECR (x-axis) and the fraction of detected molecules assigned to cells (y-axis). **(d)** ‘Barnyard’ plots showing the expression (counts) of two mutually exclusive genes, Slc17a7 (excitatory neuron marker) and Gad1 (inhibitory neuron marker) in each dataset, using either author-provided or Baysor segmentations from the same region of the cerebral cortex. **(e)** For each dataset, we assigned molecules using Baysor at ten different molecular assignment stringency thresholds. Varying this threshold enables us to calculate sensitivity (average total molecules per cell; y-axis) as a function of specificity (MECR; x-axis).

We therefore reprocessed each of the six datasets with a uniform segmentation processing pipeline. For each dataset, we applied the Baysor pipeline to an approximately 1mm x 1mm area of the cortex (a region which exhibits higher molecular counts per cell), which probabilistically assigns molecules to cells based on a joint likelihood function that considers both transcriptional composition and cellular morphology using only molecule positions^40^. This has been found to be an effective segmentation strategy in the cortex of the mouse brain, given the visible separation of molecule ‘point clouds’ marking distinct cell bodies^40^ (Figure 3a,b). By varying the threshold for assignment probability, we can vary the tradeoff between sensitivity and specificity observed for each dataset (Figure 3d,e). For example, applying a Baysor molecular assignment probability of 0.99 (most stringent) to the 10x Xenium dataset sharply reduced the degree of non-specific signal quantified by MECR, but also resulted in an approximately three-fold drop in the average number of detected molecules per cell (Figure 3e).

We leveraged this approach to generate ten sets of molecular segmentations for five of six datasets (excluding STARmap PLUS) where molecular position information was available. We used increasingly stringent molecular assignment thresholds and calculated sensitivity and specificity metrics for each set of segmentations. We emphasize that controlling for specificity enables an apples-to-apples comparison of the sensitivity of each technique that is not possible using the author-provided outputs. We find that after controlling for specificity, molecular sensitivity spans an order of magnitude between in situ technologies (Figure 3e). We observe the highest degree of sensitivity from MERSCOPE, with a nearly two-fold improvement compared to the closest technology, Xenium. This was followed closely by MERFISH and Molecular Cartography. Finally, the EEL FISH dataset exhibited relatively low sensitivity. We obtained similar results when analyzing multiple replicate slices from the same technology (Supplementary Figure 4a), or when thresholding molecular counts for any individual gene to mitigate the effects of outliers (Supplementary Figure 4b). Additionally, we observed that MERSCOPE exhibited the highest molecular counts per cell in pairwise comparisons where we only considered genes quantified by both panels (Supplementary Figure 4c-e).

Lastly, we examined how spatial location drives heterogeneity within an individual cell type, which represents a biological analysis that is particularly well-suited to imaging datasets. As a simple example we separated annotated astrocytes located within the cortex from astrocytes located within the thalamus (Figure 4a), and searched for differentially expressed genes between the two groups (results from the Xenium dataset shown Figure 4b, results from the MERSCOPE dataset shown in Supplementary Figure 5a,d). We then compared the results to scRNA-seq data, where cortical and thalamic astrocytes were separately profiled from different manually dissected samples^41^. Strikingly, when performing differential expression using the published in situ outputs, many of the top differentially expressed genes are highly expressed makers of other cell types, and are not expressed in scRNA-seq astrocytes from either the cortex or thalamus (Figure 4c).

**Figure 4.**
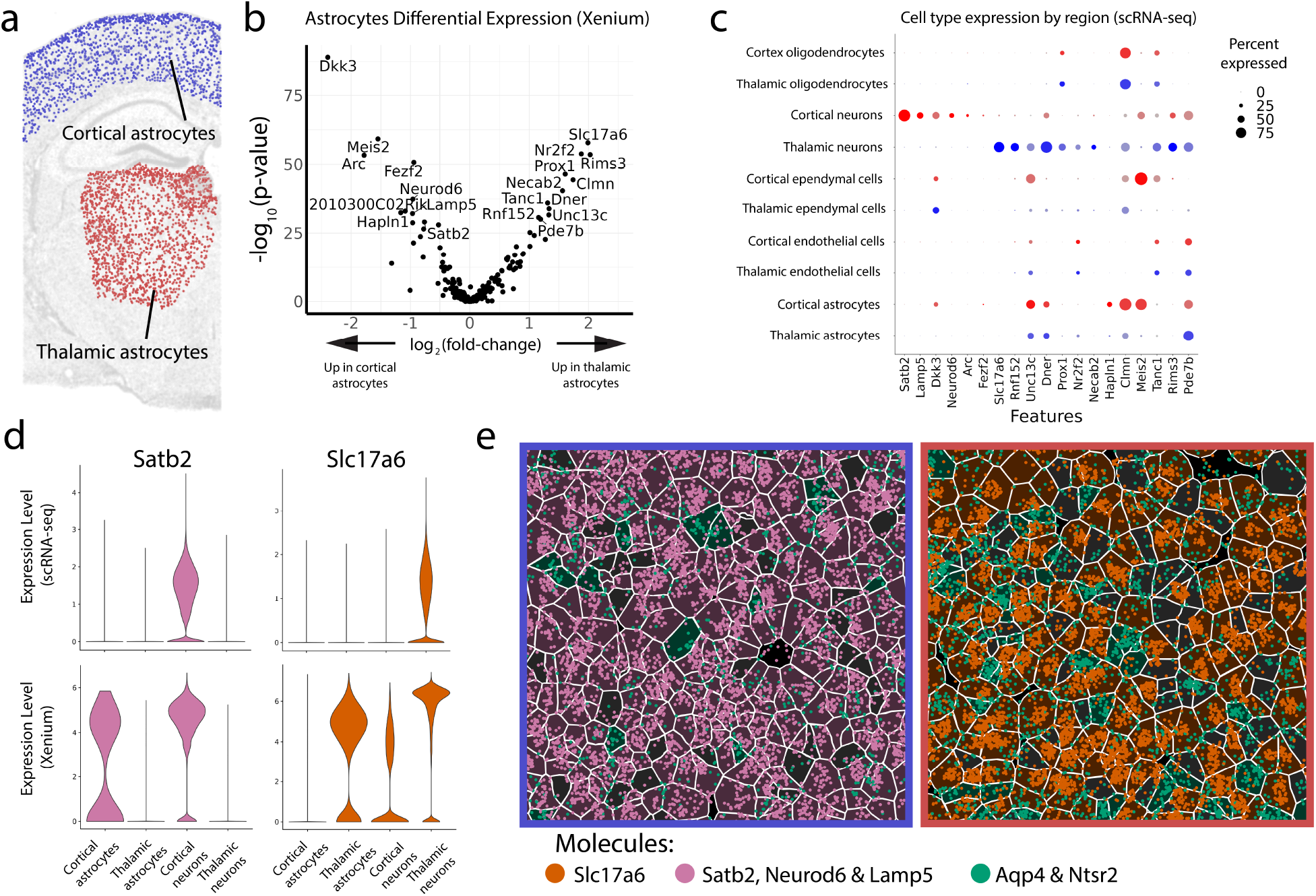
Non-specific molecular assignments confound spatial differential expression analysis. **(a)** We performed differential expression analysis between astrocytes located in the cortex (blue) and astrocytes that were located in the thalamus (red). Xenium dataset is shown. **(b)** Volcano plot of differential expression results between thalamic and cortical annotated-astrocytes in the Xenium dataset, based on author-provided segmentations. **(c)** Dot plot illustrating the expression levels of DE genes from (**b**), in a scRNA-seq dataset across cell types identified from manually dissected cortical and thalamic samples. In many cases, the identified DE genes are neuronal markers, suggesting that the DE result (obtained from astrocytes) was a result of non-specific molecular expression **(d)** Expression of Satb2 and Slc17a6 across neurons and astrocytes in the cortex and thalamus quantified by Xenium and scRNA-seq. **(e)** Zoomed view of segmentations and marker gene molecules for cortical neurons (Satb2, Neurod6 and Lamp5), thalamic neurons (Slc17a6), and pan-region astrocytes (Aqp4 and Ntsr2) in representative cortical (blue) and thalamic (red) regions. Neuronal marker molecules are mis-assigned to astrocytes, confounding differential expression analysis. The segmentation background color denotes the annotated cell type; Dark pink segmentations are cortical neurons, dark orange segmentations are thalamic neurons, dark green segmentations are astrocytes, and dark grey segmentations are assigned to one of the remaining cell types.

These results are consistent with the presence of non-specific molecular expression in in situ datasets, and the potential for a biased mis-assignment of molecules in astrocytes that belong to neighboring cells. As an example, Satb2 is a neuronal marker, and is strongly up-regulated in cortical neurons compared to thalamic neurons (Figure 4c,d). However, if a percentage of Satb2 molecules are mis-assigned to astrocytes, this gene will appear to be up-regulated in cortical astrocytes as well. Similarly, the thalamus harbors a larger population of Slc17a6+ neurons, and molecular misassignment causes Slc17a6 to be the most significantly differentially expressed gene in thalamic astrocytes (Figure 4b,e). We identified this signal in all in situ datasets (results for MERSCOPE shown in Supplementary Figure 5). Moreover, repeating analyses on the Baysor-reprocessed datasets alleviated, but did not eliminate, this non-specific signal (Supplementary Figure 5). These results highlight the danger of non-specific molecular signals not only for comparative benchmarking, but also for downstream molecular analyses. We encourage users to carefully consider this factor when interpreting differential expression analyses in situ gene expression datasets.

## Discussion

In this study, we performed a comparative benchmarking analysis of six multiplexed in situ gene expression profiling datasets spanning both academic and commercial technologies. In each case, we utilize datasets generated by the technology’s inventors, aiming to compare datasets that demonstrate the full capacity of each method. While our initial analyses focus on author-generated outputs, we also re-analyze five of the six technologies using a common segmentation pipeline, which better enables us to compare sensitivity and specificity across datasets.

While all technologies generate useful datasets and faithfully recover the spatial organization of mouse brain tissue, we find that Vizgen’s MERSCOPE datasets exhibit the best performance. This is clearly reflected in our technical metrics computed on both the MERSCOPE published outputs and following re-segmentation, where MERSCOPE exhibits the optimal trade-off between sensitivity and specificity. Moreover, MERSCOPE achieves this performance while also featuring a large panel size of 483 genes. Our analyses find that the Xenium technology, which featured a panel of 247 genes, also exhibited good performance, but we also find that the segmentations returned by the Xenium computational pipeline were far larger and more lenient than what we observed in other technologies. These segmentations increase the number of molecules detected per cell, at the expense of a substantial increase of incorrect molecular assignment.

We observed that the Resolve Molecular Cartography dataset and MERFISH datasets exhibited similar sensitivity, albeit with very different panel sizes. The Molecular Cartography dataset contained just 99 genes, demonstrating impressive sensitivity on a per-gene basis. In contrast, the MERFISH dataset featured the largest panel (1147 genes), demonstrating the ability to achieve strong performance while maintaining a high degree of plexity. The EEL FISH dataset was the least sensitive of all technologies profiled. The reduced sensitivity in EEL FISH is a well-described tradeoff of the technology^9^, which emphasizes fast imaging time and minimized autofluorescence by electrophoretically transferring mRNA transcripts to a secondary slide.

Our study has a few limitations. First, our focus on mouse brain enabled us to compare a broader set of both commercial and academic technologies for which publicly available datasets are available. While this is a commonly used tissue for evaluating spatial analyses platforms, it represents only a single biological context. Moreover, our analyses focused on fresh-frozen tissue, representing a complementary evaluation to two recent studies which specifically evaluated performance on FFPE tissues^33,34^. Finally, our Baysor analysis focused on cells in the mouse cortex, and as with all in situ benchmarking analyses, may be influenced by the specific composition of target genes and probes in the panel. More broadly however, our findings highlight the importance of considering both sensitivity and specificity when evaluating the performance of in situ technologies, and we believe this will be an important consideration across tissue types.

A substantial component of non-specific signal in imaging-based spatial transcriptomic datasets appears to be largely driven by misassignment of molecules from neighboring cells. We anticipate the need for two types of computational methods that can address this issue. First, methods for improved segmentation - and in particular those that use both nuclear and membrane stains as well as the position and identity of detected molecules in 3-dimensional space - will improve the overall results. However, no segmentation algorithm is perfect, and segmentation can be especially challenging when cells are densely packed or have complex morphology^28,40^. We therefore anticipate that new computational methods, analogous to those that control for ambient RNA in scRNA-seq^38,42^, will be able to account for and correct for non-specific biases that arise in downstream analyses. As the field of spatial transcriptomics continues to develop, we hope that our analysis provides a useful and interpretable framework to assess new technologies, datasets, computational methods, and tissue preparation techniques going forward.

## ACKNOWLEDGEMENTS

We thank members of the Satija Lab (and in particular, Paul Hoffman), Jean Fan, and Brendan Miller for helpful discussions. This work was supported by the Chan Zuckerberg Initiative (EOSS5-0000000381, HCA-A-1704-01895 to R.S.), and the National Institutes of Health (RM1HG011014-02 and 1OT2OD033760-01 to R.S).

## CONFLICT OF INTEREST STATEMENT

A.H. was employed by 10x Genomics from July 2020 to September 2021 and owns stock in the company. In the past 3 years, R.S. has received compensation from Bristol-Myers Squibb, ImmunAI, Resolve Biosciences, Nanostring, 10x Genomics, Neptune Bio, and the NYC Pandemic Response Lab. R.S. is a co-founder and equity holder of Neptune Bio.

## DATA AND CODE AVAILABILITY

Code to generate plots in the main and supplementary figures via a reproducible snakemake workflow accessible at https://github.com/AustinHartman/spatialbenchmark_reproducibility. Dataset download is described in the “Data acquisition” section of the methods.

## SUPPLEMENTARY METHODS

### Data acquisition

Spatial transcriptomic and scRNA-seq datasets used in this manuscript can be freely downloaded from different public portals and company websites. We finalized our list of datasets to include based on what was publicly available on June 1st, 2023, and may update this manuscript in the future to include mouse brain datasets that have been released since that time. Instructions to download datasets and load as Seurat objects are summarized for each dataset below:

*(Xenium; 10x Genomics)*: The Xenium dataset represents a fresh frozen mouse brain coronal section, with a 248-gene panel. We downloaded the dataset from the 10x Genomics website at *https://www.10xgenomics.com/products/xenium-in_situ/mouse-brain-dataset-explorer*.

Downloaded files included molecular identity and coordinate information, cell segmentation boundary coordinates, a quantified single-cell gene expression matrix, and the spatial coordinates of each cell centroid. The outputs were generated using “Xenium Onboard Analysis” software version 1.0.2. The LoadXenium function in Seurat uses some of these files to construct a Seurat object.

(*MERSCOPE; Vizgen)*: The MERSCOPE dataset, called the “Vizgen MERFISH Mouse Receptor Map”, uses of a 483-gene panel. Three full coronal slices, with three replicates each are available for download. In this paper, slice 2, is used and is accessible from the Vizgen website at https://info.vizgen.com/mouse-brain-data. Downloaded files include stain images for segmentation, a cell by gene matrix, cell segmentation boundary coordinates, and molecular identity and coordinate information. The LoadVizgen function in Seurat uses some of these files to construct a Seurat object.

(*Molecular Cartography, Resolve Biosciences*): The Resolve Biosciences dataset is available on request, which is submitted on the Resolve Biosciences website at https://resolvebiosciences.com/open-dataset/?dataset=mouse-brain-2021. The downloadable files include DAPI stains, a quantified single-cell gene expression matrix, detected molecule positions and identities, and segmentation coordinates for each cell provided in separate files. The LoadResolve function, which is defined in the code provided with this manuscript, uses some of these files to construct a Seurat object.

(*MERFISH)*: The complete MERFISH dataset quantifies 1147 genes and over 10 million cells in the mouse brain^13^. Numerous coronal and sagittal sections spanning the length of the brain are available and registered using the Allen CCFv3^43^. Data is available to download from the Brain Image Library here: https://download.brainimagelibrary.org/29/3c/293cc39ceea87f6d/. Raw and normalized counts for individual mice are provided in separate H5AD files containing a quantified single-cell gene expression matrix, cell center information, FOV information, and slice and sample identification information. Sample index 21 and 22 from “raw_counts_mouse2_coronal.h5ad” were used in this paper. The LoadMERFISH23 function, which is defined in the code provided with this manuscript, uses the H5AD output to construct a Seurat object.

(*STARmap PLUS*): The STARmap PLUS dataset contains 17 coronal and 3 sagittal mouse brain sections but somewhat limited output files. Available outputs are a quantified single-cell gene expression matrix and the position of each cell. Position and identity of detected molecules are not provided which is required for molecule assignment using Baysor. The dataset is downloadable from the Broad Single Cell Portal at *https://singlecell.broadinstitute.org/single_cell/study/SCP1830/spatial-atlas-of-molecular-cell-types-and-aav-accessibility-across-the-whole-mouse-brain*. The LoadSTARmapPlus function, which is defined in the code provided with this manuscript, uses the H5AD output to construct a Seurat object.

(*EEL FISH*): The EEL FISH dataset assays a full sagittal section and is downloadable from http://mousebrain.org/adult/. The files include positions and identities of all detected molecules in CSV and parquet format, and a gene expression matrix and cell positions stored in H5AD format. The LoadEELFISH function, which is defined in the code provided with this manuscript, uses the H5AD output to construct a Seurat object.

(*Allen Brain Cortex Reference*): This is a scRNA-seq dataset generated with the SMART-seq2 protocol^44^. The dataset contains roughly 14,000 cells from the cortex of an adult mouse and a well-annotated cortical taxonomy^45^. Data is available for download from https://www.dropbox.com/s/cuowvm4vrf65pvq/allen_cortex.rds.

(*Linnarsson Adolescent Mouse Nervous System Reference*): This is an scRNA-seq dataset generated with 10x Genomics Chromium Single Cell 3’ version 1 and 2 kits. Data is available for download from http://mousebrain.org/adolescent/downloads.

### Replicate correlation

The replicate plots display the total number of detected molecules for each probe (gene targeting and negative targeting) for two replicates. The MERSCOPE plot shows replicates one and two from slice two. The Xenium plots show replicates one and two from the fresh frozen mouse brain public dataset. STARmap PLUS displays sagittal section 2 and sagittal section 3. MERFISH shows sample index 22 and sample index 21 from mouse 2 coronal sections. Replicates for Molecular Cartography and EEL FISH are not shown because only one sample is available. For more details on the specific datasets see the “Data acquisition” section of the methods.

### Preprocessing, clustering, and annotation

*Quality control:* For each dataset, we used the author-provided gene expression matrix to select cells, and did not perform any additional quality control or filtering steps. We note that while filtering steps are common in scRNA-seq analysis, for example to remove material originating from empty droplets, filtering cells in spatial analysis would generate gaps in the tissue.

*Normalization, dimensional reduction, and cell annotation*: We performed standard log-normalization for each dataset using the NormalizeData function in Seurat v4.3. We reduced the dimensionality of each dataset using principal components analysis (RunPCA), using default parameters and including all targeted genes. We further reduced dimensionality to two dimensions using the RunUMAP function, to generate the visualizations in Figure 1d.

We performed cell annotation using an scRNA-seq (10x Genomics Chromium system) dataset representing a comprehensive survey of 133 manually dissected anatomical regions from the adolescent mouse nervous system^36^. For this dataset, we retained cells expressing between 200-3,000 unique features, and with less than 5% of molecules derived from mitochondrial transcripts. We also removed any samples from which less than 500 cells were obtained. We constructed a reference from this dataset using the standard Seurat scRNA-seq including the NormalizeData, FindVariableFeatures (with 4,000 variable features), ScaleData, and RunPCA (using 100 principal components) functions.

We used the Seurat label transfer workflow^46,47^, including the FindTransferAnchors (using one non-default parameter; reduction=‘cca’) and MapQuery functions, to transfer scRNA-seq derived annotations to each of the in situ datasets. We transferred both broad class labels (Fig. 1a, d) from the “Class” metadata column, as well as the higher-resolution “Taxonomy Group” labels (Supplementary Figure 3a). For downstream analyses in this study, we use the broad ‘Class’ cell type labels, as each technology includes sufficient gene panels and data quality to confidently discriminate cells between these groups. We verified the accuracy of our transferred annotations by performing differential expression (Wilcoxon test implemented using the Presto package^48^) on each in situ dataset based on the annotated labels, and visualizing the top marker genes (Figure 2a).

### Specificity metrics

Specificity in single-cell sequencing datasets is often assessed using a ‘barnyard’ analysis^39^, where mixtures of cells from different species are profiled together in a single sample. A bona fide single cell should never co-express transcripts from multiple species, enabling the detection of doublets or non-specific molecular assignments. While the in situ datasets assessed here all contained only mouse cells, we reasoned that we could perform an analogous analysis by testing for mutual expression of markers of multiple distinct cell types (which should not be co-expressed). We found that in scRNA-seq data, marker genes of different cell types exhibited mutually exclusive expression, indicative of high levels of specificity (Supplementary Figure 2b).

We first identified a set of marker genes for each of 7 broad cell types from the Linnarsson scRNA-seq dataset. Cell type markers are identified for all cell types using the Linnarsson reference dataset with a Wilcoxon test implemented by Presto^48^. Cell type markers in the Linnarsson scRNA-seq dataset are filtered based on two criteria: they must exhibit more than 25% non-zero expression in the marker population, and less than 1% non-zero expression in all other cells. Any gene that was identified as a marker for multiple cell types was removed. From this final marker set, we generated a list of ‘mutually exclusive’ gene pairs.

For each in situ dataset, we considered only the mutually exclusive gene pairs where both members were present in the target gene panel. We computed the mutually exclusive co-expression rate (MECR) for each gene pair, by dividing the fraction of cells which express both markers by the fraction of cells which express at least one of the markers. Boxplots showing the range of MECR values across gene pairs are shown in Figure 2e. We used the average MECR value across all gene pairs as an estimation of MECR for the dataset (as in Figure 3c, e).

### Baysor segmentation

We used Baysor v0.6.1^40^,and leveraged the ability to probabilistically assign molecules to cells using only the positions of detected molecules. Since the computational requirements for running Baysor scale with the total number of molecules, we subsetted each dataset to a roughly 1,000 by 1,000 micron region. We excluded the STARmap PLUS dataset since molecule positions were not publicly available.

Baysor is provided with three inputs; a CSV containing molecule positions and identities, ‘scale’ and ‘min-molecules-per-cell’. ‘Scale’ is the expected cell radius which was set for each to represent an expected cell radius of 6 microns, though the parameter value we used varied depending on the zoom level and coordinate system for the provided images: Xenium (6), MERSCOPE (6), MERFISH (6), Molecular Cartography (30), EEL FISH (60). The ‘min-molecules-per-cell’ parameter is the number of molecules required for a cell to be called. We set this to 100, but reduced it to 25 for EEL FISH based on the lower sensitivity.

Baysor assigns a confidence score to each molecule allocated to a cell, with the score ranging between 0 and 1. We compute multiple gene expression matrices with ten different assignment cutoffs (0, 0.25, 0.5, 0.55, 0.6, 0.65, 0.7, 0.75, 0.8, 0.85, 0.9, 0.95, 0.97, 0.98, and 0.99), in order to to vary the specificity of segmentations. We then compute specificity metrics (i.e. MECR) and sensitivity metrics (i.e. average number of detected molecules per cell) for each of these segmentations to generate the curves in Figure 3e.

---

**Supplementary Figure 1.**
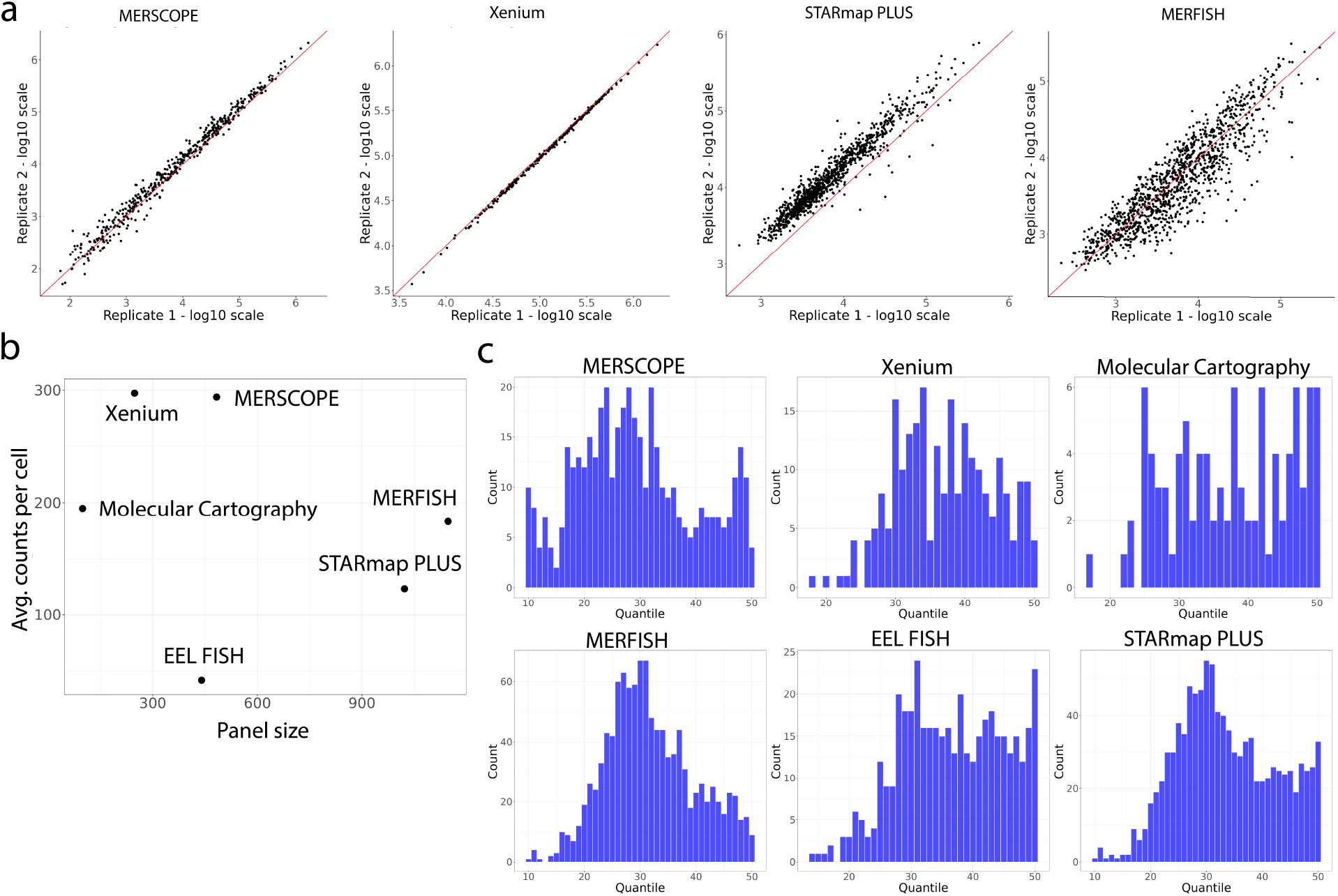
**(a)** Log-transformed pseudobulk expression for all genes. Plots are shown for technologies where replicates (i.e. two adjacent tissue slices) are available. **(b)** The average molecular counts detected per cell (using author-provided segmentations) does not show a relationship with the total number of genes targeted. **(c)** Distribution of gene expression quantiles, derived from the Linnarsson scRNA-seq dataset, for each gene which is targeted in each panel.

**Supplementary Figure 2.**
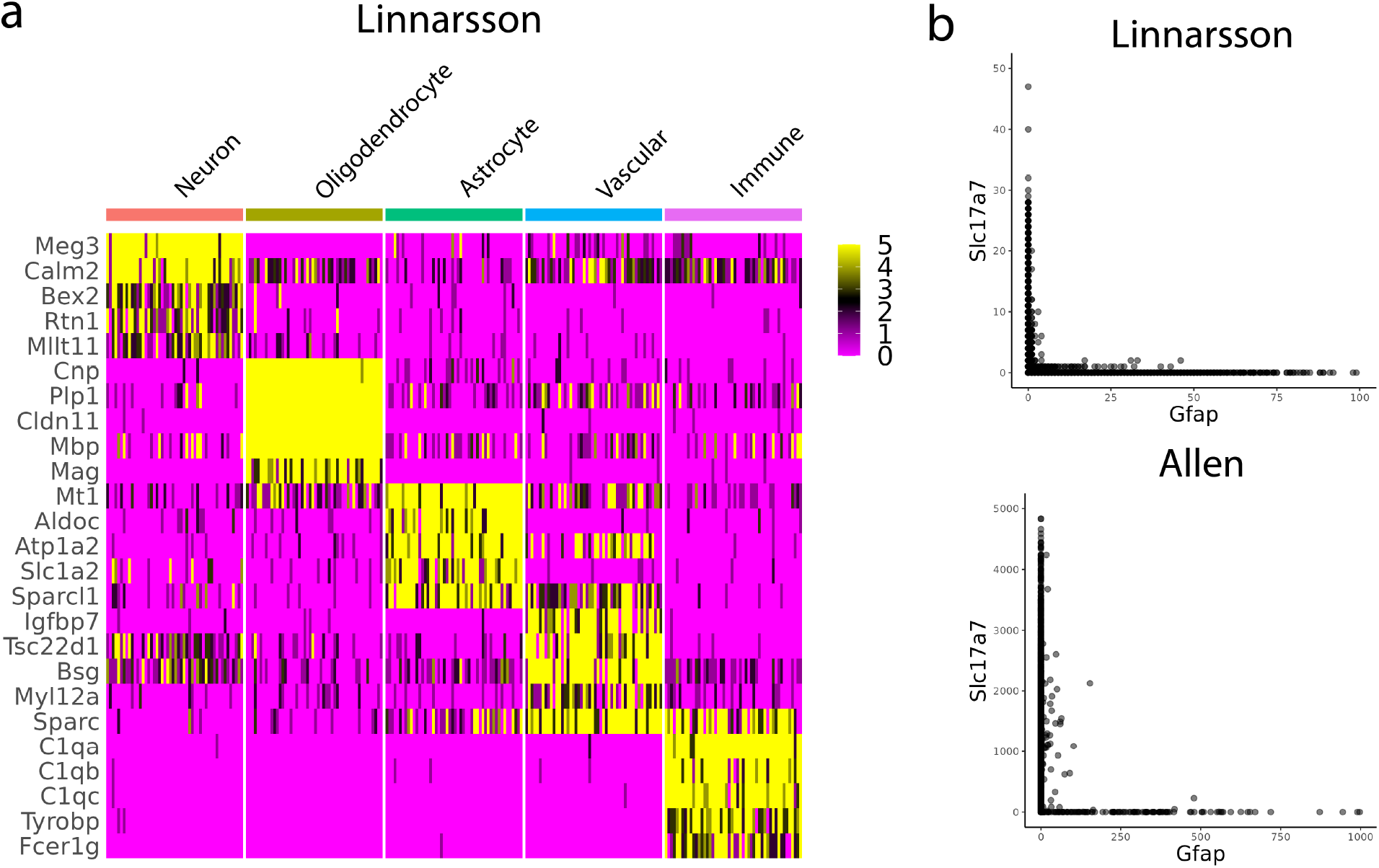
**(a)** Heatmap displaying gene expression of marker genes for 250 total sampled cells from 5 major cell types. Same as Figure 2a but for the Linnarsson scRNA-seq dataset. Raw counts thresholded at 5 are displayed. **(b)** Slc17a7 and Gfap counts in cells from the Linnarsson Reference and Allen cortex dataset are shown. Both are scRNA-seq datasets, and in contrast to in situ technologies (Figure 2c), exhibit minimal evidence of mutual co-expression of these markers.

**Supplementary Figure 3.**
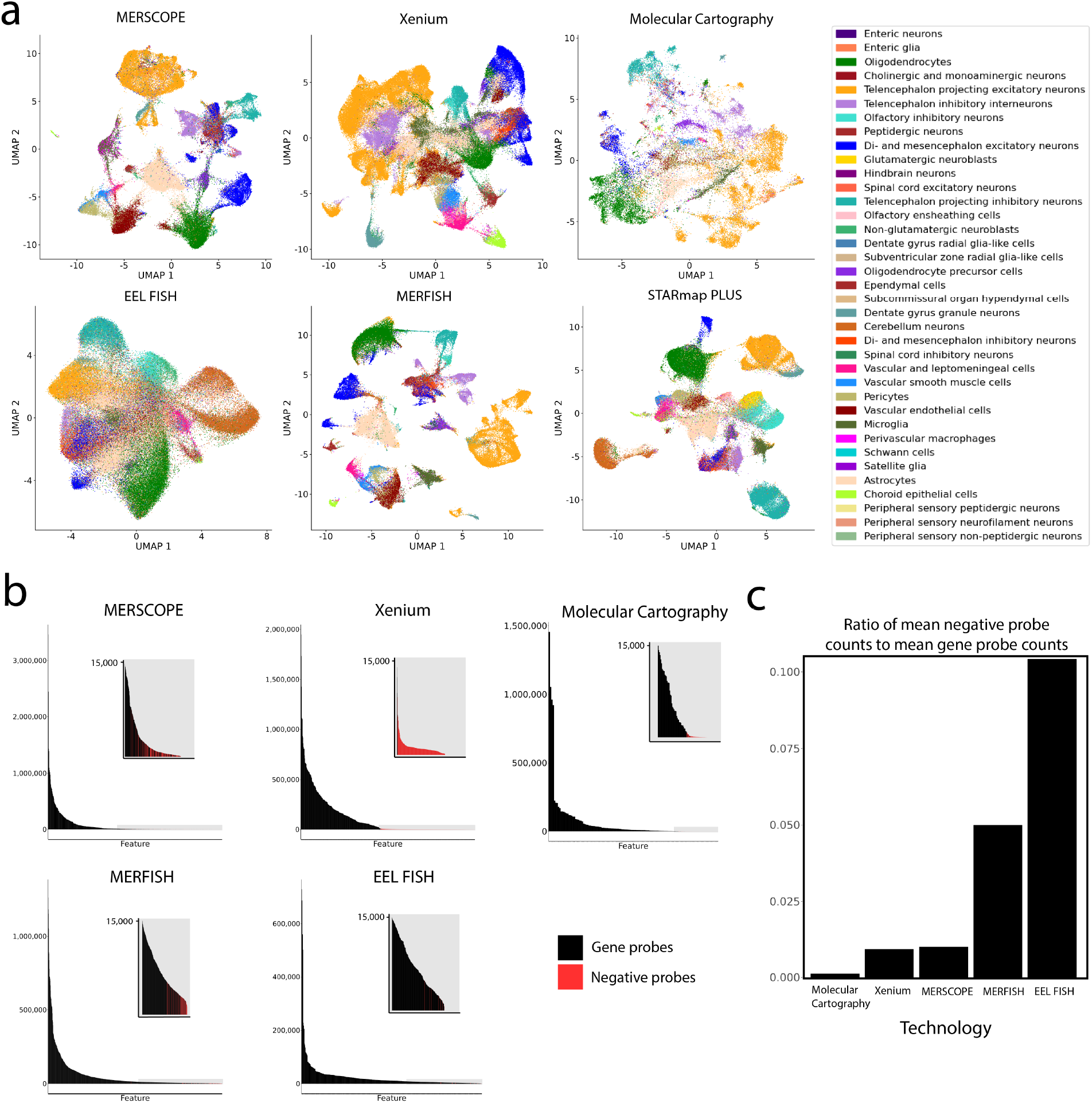
**(a)** UMAP visualization of each dataset. Same as in Figure 1d, but cells are colored by higher resolution annotations, derived from scRNA-seq label transfer. **(b)** A rank-sorted comparison of the expression levels of gene targeting probes (in black) and negative targeting control probes (in red). A truncated subplot is provided to emphasize probes at lower expression levels, with the relevant sections of the rank plot highlighted in grey. **(c)** The ratio of the mean counts of negative targeting probes to gene targeting probes. A ratio value of 0.1 signifies that for every detected gene probe, there are 0.1 negative probes detected.

**Supplementary Figure 4.**
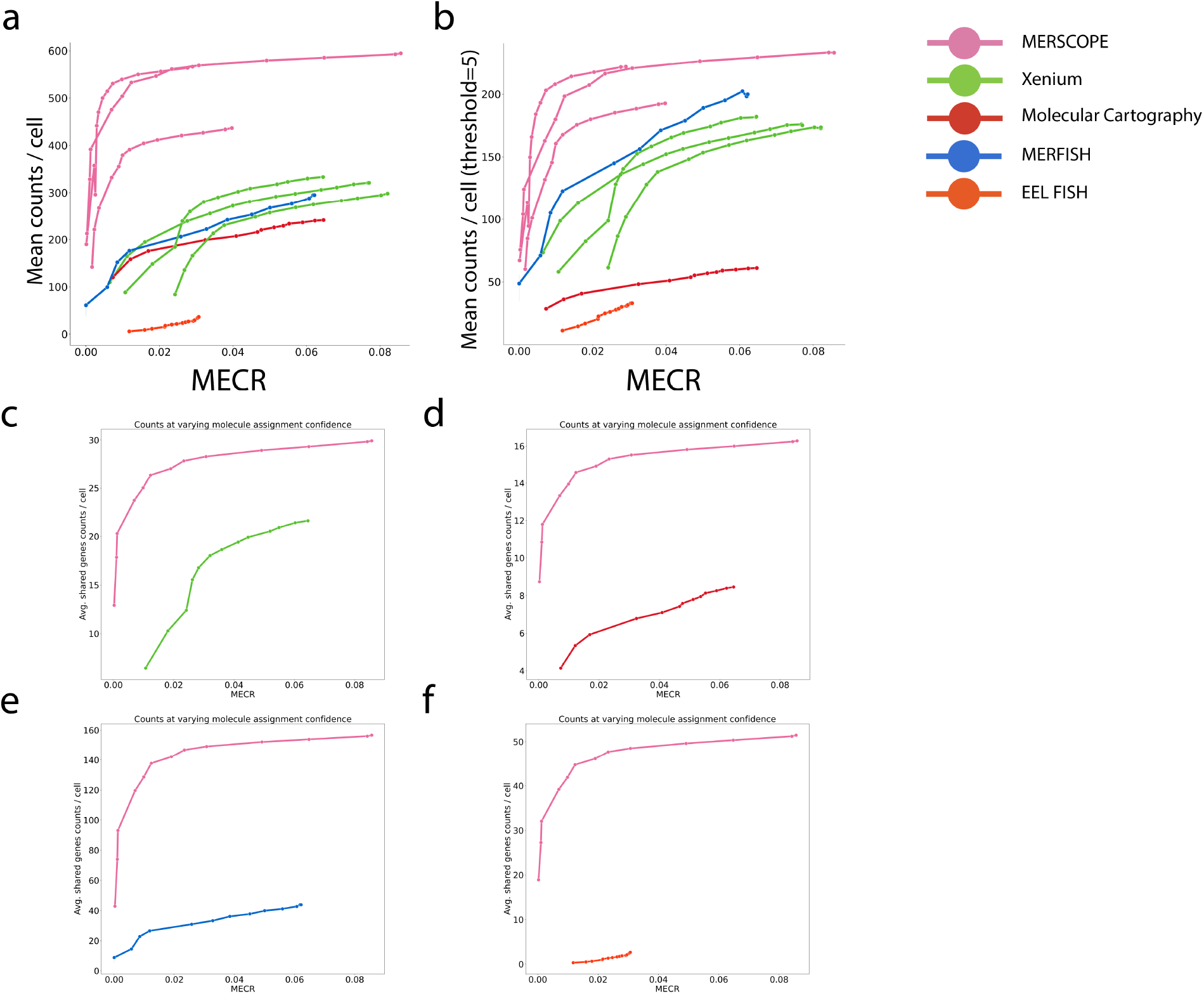
**(a)** Sensitivity versus specificity curves shown for multiple experimental replicates. Same as Figure 3e, but multiple replicates are shown when available for the same technology **(b),** Same as panel **(a)**, but the mean is computed after thresholding the maximum counts for each gene to 5, to reduce the effect of highly expressed outliers, and limiting the effect that a single gene can have on this metric. Technologies that use large panels, like MERFISH, exhibit a more significant relative improvement when compared to the absence of thresholding, as opposed to technologies that use small panels, such as Molecular Cartography. **(c)** Mean counts for genes targeted by both MERSCOPE and Xenium versus MECR. There are 27 genes targeted by both panels. **(d)** Same as **(c)**, but Molecular Cartography shown instead of Xenium. There are 16 genes targeted by both panels. **(e)** Same as **(c)**, but MERFISH shown instead of Xenium. There are 142 genes targeted by both panels. **(f)** Same as **(c)**, but EEL FISH shown instead of Xenium. There are 53 genes targeted by both panels. For panels **(c)** through **(e)** Slc17a7 is excluded because it is an extremely highly detected outlier gene.

**Supplementary Figure 5.**
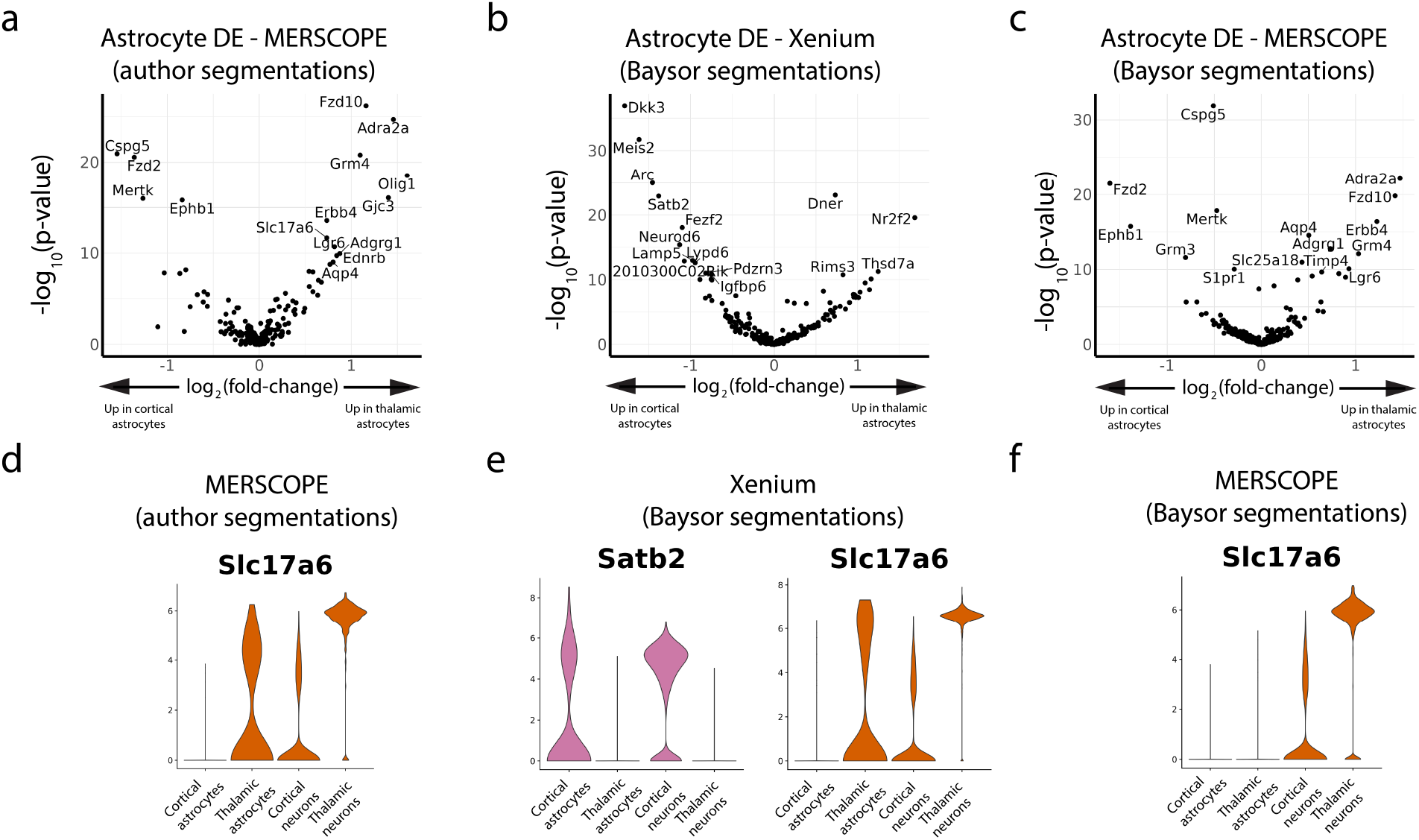
**(a)** Volcano plot displaying differentially expressed genes between cortical astrocytes and thalamic astrocytes in MERSCOPE dataset using author-provided segmentations. **(b)** Volcano plot displaying differentially expressed genes between cortical astrocytes and thalamic astrocytes in Xenium dataset using Baysor segmentations. **(c)** Volcano plot displaying differentially expressed genes between cortical astrocytes and thalamic astrocytes in MERSCOPE dataset using Baysor segmentations. **(d)** Expression of Slc17a6 in neurons and astrocytes from the thalamus and cortex in the MERSCOPE dataset using author-provided segmentations. **(e)** Expression of Satb2 and Slc17a6 in neurons and astrocytes from the thalamus and cortex in then Xenium dataset using Baysor segmentations. **(f)** Expression of Slc17a6 in neurons and astrocytes from the thalamus or cortex in MERSCOPE dataset using Baysor segmentations. Satb2 is not shown in **(d)** and **(f)** because it is not included in the MERSCOPE panel.

## Notes

### Summary of Updates

Supplemental figures updated.

